# Fine-scale behavioural dynamics separates adaptive sickness behaviour from injury and infection pathology

**DOI:** 10.64898/2026.04.21.719817

**Authors:** Alejandro V. Cano, Dhobasheni Newman, Katy M. Monteith, Vasilis Dakos, Pedro F. Vale

## Abstract

Animals commonly reduce activity during infection, but it remains unclear when such “sickness behaviours” reflect adaptive host regulation versus injury responses or pathogen-driven pathology. We combined high-resolution behavioural phenotyping with experimental partitioning of injury, immune stimulation, and live bacterial infection to dissect these effects in *Drosophila melanogaster*. Using an outbred fly population, we tracked individual locomotor activity at 1 minute resolution for 400 hours in males and females subjected to one of four treatments: unhandled control, septic injury with buffer, immune stimulation with heat killed *Pseudomonas entomophila*, or septic infection with live *P. entomophila*. Across the early post-challenge window, total activity diverged most strongly between the unhandled control and the challenged treatments. Fine-scale analyses revealed that immune challenge primarily reshaped the microstructure of movement: the number of activity bouts changed little, whereas active bout lengths were shorter and inactive periods were modestly prolonged, especially in females. Among infected flies, individuals that died early exhibited a collapse of bout structure, showing prolonged inactivity punctuated by brief, sometimes intense, movements, whereas longer-lived infected flies maintained patterns closer to controls. These behavioural differences were detectable in the very early stages of infection: within the first 24-hours post-infection early-dying flies had both a reduced probability of initiating movement from rest and a reduced probability of sustaining on-going activity, consistent with pathogen-induced debilitation; longer-lived infected flies showed near-normal state-transition profiles. This early suppression of activity in acutely dying males was positively associated with subsequent survival time, suggesting a contribution of adaptive restraint alongside pathology. By resolving fine-scale behaviour and state-transition dynamics, our approach clarifies which aspects of “sickness behaviour” are likely adaptive and which signal pathology.

## INTRODUCTION

Animals commonly exhibit coordinated changes in behaviour during infection, including reduced locomotion, altered feeding, increased sleep or rest, and changes in social interaction^1–4^. Collectively termed sickness behaviours, these responses are phylogenetically widespread and have been documented in both vertebrates and invertebrates^3–5^. In vertebrates, the syndrome is classically mediated by immune-to-brain signalling, with cytokines and prostaglandins acting on neural circuits to reduce activity, induce lethargy, suppress appetite, and increase sleep^2,3^. In invertebrates, immune activation and infection likewise modulate behaviour, though the mechanistic routes can differ and often involve direct integration of immune and neuroendocrine pathways^6–9^. The ubiquity of these behavioural changes across taxa has prompted the influential hypothesis that sickness behaviours are adaptive host strategies that reallocate energy to immune defence, reduce exposure to further risk, and potentially limit pathogen transmission^10–12^.

Despite their apparent generality, establishing the adaptive value of sickness behaviours in any given host-pathogen system remains challenging. While there is compelling evidence for host-driven behavioural changes in response to perceived (but not necessarily live) infection^4,6,13,14^, behaviours attributed to “sickness” during live infections can arise from multiple, non-mutually exclusive processes: an active host response (e.g., metabolic reprogramming for more efficient immune responses^15–17^), nonspecific responses to tissue damage (e.g., injury-induced wound repair responses^18,19^), and direct pathogenesis caused by the pathogen^20–24^. Disentangling these contributions empirically requires designs that partition the effects of injury, sterile immune stimulation, and live infection, while quantifying behaviour with sufficient temporal resolution to reveal both acute and longer-term dynamics.

*Drosophila melanogaster* is an experimentally tractable model of both infection and behaviour^25–27^, providing a powerful system with which to address these questions. We focus here on bacterial infection with *Pseudomonas entomophila*, an established Drosophila pathogen that elicits a strong antibacterial response and can cause substantial pathology^28–30^. Although widely used for enteric infection, *P. entomophila* also induces systemic immune activation and morbidity after septic injury^31,32^, providing a tractable model of acute bacterial challenge in which both immune stimulation and pathogen-derived pathology can shape behaviour. In parallel, Drosophila locomotor activity is readily quantified at high temporal resolution^33–36^. Immune activation and infection can modulate these outputs: in some cases, immune challenge can increase sleep or reduce activity in flies^37,38^, which is consistent with reallocation of resources toward defence, while in other contexts, immune-mediated increases in activity have also been detected^39^.

Activity is a particularly informative readout for parsing these effects in Drosophila for several reasons. First, locomotor activity is energetically costly and tightly coupled to metabolic state^35,40,41^; reductions in movement are therefore expected if hosts reallocate limited resources from performance to defence and repair. Second, activity integrates multiple dimensions of sickness behaviour, including sleep regulation, arousal, and motivation^33,42^, which allows inference about both the initiation and maintenance of movement. Third, high-throughput monitoring across days captures the interaction between acute responses and longer-term survival, and enables analysis of within-day behavioural structure, which can reveal whether infection alters the probability of starting, sustaining, or terminating movement. Finally, flies show sexual dimorphism in immune and behavioural responses^38,43,44^, motivating explicit tests for sex-specific sickness behaviours.

In this study, we combine these advantages to quantify how injury, immune stimulation, and live bacterial infection shape locomotor behaviour in male and female *D. melanogaster*. Using an outbred laboratory population, we recorded individual activity continuously, at 1-minute resolution, for 400 hours (*∼*16 days) following one of four treatments (Fig. 1A): unhandled control (naive) or pricked with a sterile buffer (injury), heat-killed *P. entomophila* (immune stimulation), or live *P. entomophila* (infection). We analysed fly behavioural responses at multiple scales. At the coarse scale, we assessed treatment effects on total activity and survival. At a finer scale, we characterized the organisation of active and inactive bouts over time. We further modelled the minute-to-minute state switching to estimate the probabilities of initiating versus sustaining movement. Finally, because survival rates are heterogeneous between individuals, we stratified live-infected flies by survival time to test whether acutely dying individuals exhibit distinct patterns of activity compared to chronic survivors. By partitioning the drivers of behavioural change and resolving the fine-scale structure of movement, our approach clarifies when and how reduced activity reflects adaptive host response versus injury- or pathology-driven impairment.

**Figure 1:**
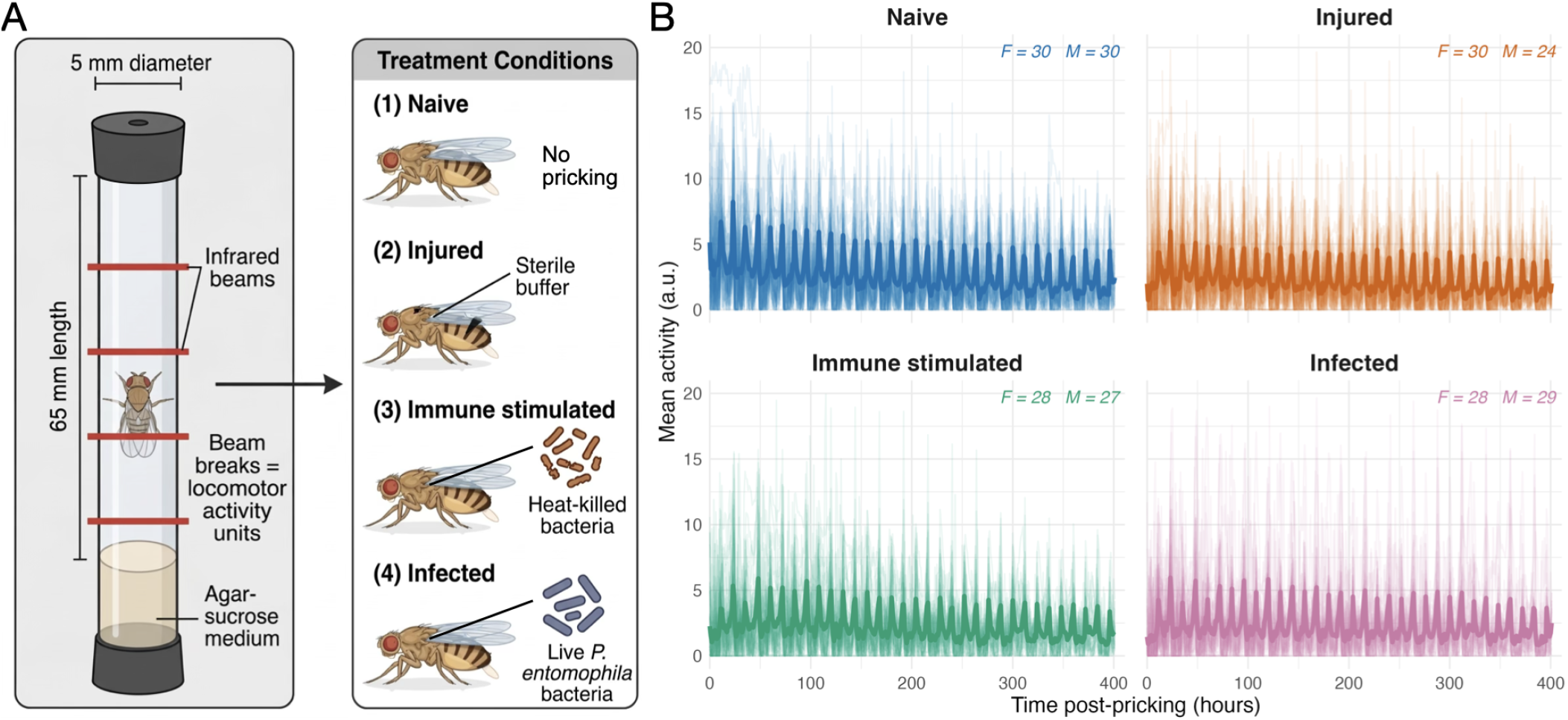
Experimental setup and locomotor activity across immune challenge treatments. **(A)** Schematic representation of experimental setup. **(B)** Individual fly activity timeseries (faded lines) and treatment mean (bold line) plotted as mean activity (number of beam breaks per minute) over 400 hours post-infection for each treatment group (naive, injured, immune stimulated, injured). Sample sizes per sex are annotated in each panel (F = females, M= males). Treatment: naive (blue), injured (orange), immune stimulated (heat-killed Pe - green), infected (live Pe – pink).

## RESULTS

### Bacterial infection reduces locomotor activity in a sex-specific manner

All four treatment groups displayed persistent oscillations in locomotor activity throughout the experiment, alternating between periods of high and low activity (Fig. 1B). This rhythmic pattern was maintained across the full 400-hour recording window, showing that injury and immune challenges do not completely disrupt activity patterns. A comparison of daily mean activity across treatments revealed that naive flies maintained the highest mean activity levels (Fig. 2A and Fig. S1). Over the course of the 400-hour recording period, activity trajectories diverged over time between treatments in a sex-dependent manner (significant three-way Treatment × Sex × Hour interaction: *F*_3,69023_ = 14.99, *p* < 0.001). All flies experienced significant mortality throughout the experiment (Fig. 2B), which is expected given the deteriorating environment experienced within a DAM tube^45,46^. Kaplan–Meier curves suggest that *P. entomophila*-infected flies experienced the steepest decline in survival fraction, while naive flies exhibited the highest survival ((Fig. 2B)). However, we did not detect any statistically significant differences in mortality risk between any treatment pair (Cox-PH *χ*^2^ = 5.42, *df* = 4, *p* = 0.2; all BH-corrected pairwise comparisons *p >* 0.62). The absence of statistically significant survival differences between treatments suggests that the activity differences we observe are unlikely to be driven by selective mortality of the least active individuals. Rather, they likely reflect a behavioural response to injury and immune challenge.

**Figure 2:**
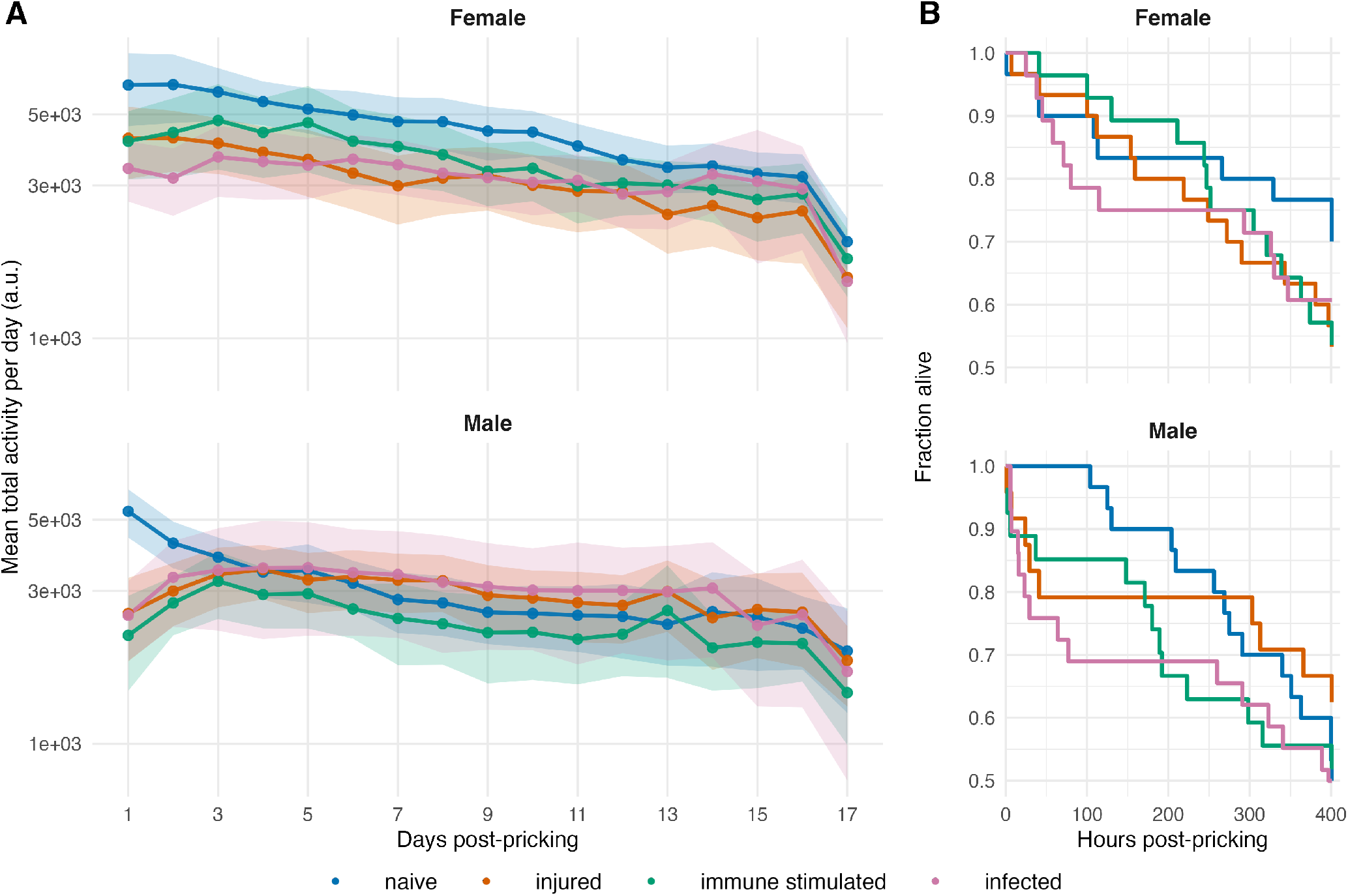
Locomotor activity and survival across immune challenge treatments. **(A)** Daily mean locomotor activity across treatments and sexes. Mean total activity per day (arbitrary units, log scale) for each treatment group over 17 days post-injection, shown separately for females (top) and males (bottom). Points represent daily means; shaded ribbons indicate 95% confidence intervals. Colours: naive (blue), injured (orange), immune stimulated (green), live *P. entomophila* (pink). **(B)** Survival curves showing the fraction of flies alive over time, plotted separately for females (top) and males (bottom). Colours correspond to treatment groups as in panel A. Treatments: naive (blue), injured (orange), immune stimulated (heat-killed *Pe* — green), infected (live *Pe* — pink).

Given that the largest difference in activity between naive controls and all other challenged treatments was observed within the first 3 days (Fig. 2A and Fig. S1), and that considerable mortality occurred within the same period (Fig. 2B), we decided to focus on the first three days to characterize early treatment effects in detail. Within this window, naive flies maintained significantly higher mean activity than all challenged groups, with the effect being most pronounced and consistent in females (naive vs. injury: Δ = 1.41, *p* = 0.0035; naive vs. immune stimulation: Δ = 1.23, *p* = 0.019; naive vs. infection: Δ = 1.97, *p* < 0.0001; Tukey-corrected pairwise contrasts from LMM). In males, the same directional trend was observed but was weaker and only marginally significant for injury and infection comparisons (*p* = 0.051 and 0.054, respectively), with the immune stimulation treatment showing significantly lower activity compared to naive flies (Δ = 1.21, *p* = 0.024). No significant differences in activity were detected among the three challenged treatments in either sex during this period (all *p* > 0.30; Fig. 2A and Fig. S1).

Overall, these results show that immune challenge suppresses locomotor activity over time in a treatment- and sex-dependent manner, with the strongest and most consistent effects in females during the first three days post-pricking. To understand the basis of this suppression, we next examined whether it reflects an overall reduction in movement or a restructuring of the temporal organization of activity itself.

### Immune challenge restructures locomotor activity during the acute response time window

To characterise how immune challenge affects the fine-scale temporal structure of locomotion, we analysed the organisation of active and inactive bouts within 6-hour windows over the first three days. Across all four treatments, the number of active bouts per 6-hour window was broadly similar throughout the first three days post-pricking, with no significant effect of treatment on bout frequency (LMM: *F*_3,206_ = 1.60, *p* = 0.190; Fig. 3A). naive females produced a median of 28 active bouts per 6-hour window compared to 27–34 for challenge groups, and males showed a similar pattern (19–22.5 bouts per window). Females produced significantly more active bouts than males on average (Δ = 8.4 bouts, *SE* = 1.22, *p* < 0.001), but this difference was consistent across all treatment groups (Treatment × Sex interaction: *F*_3,206_ = 1.72, *p* = 0.165), indicating a treatment-independent baseline sex difference in bout frequency rather than a differential response to immune challenge.

**Figure 3:**
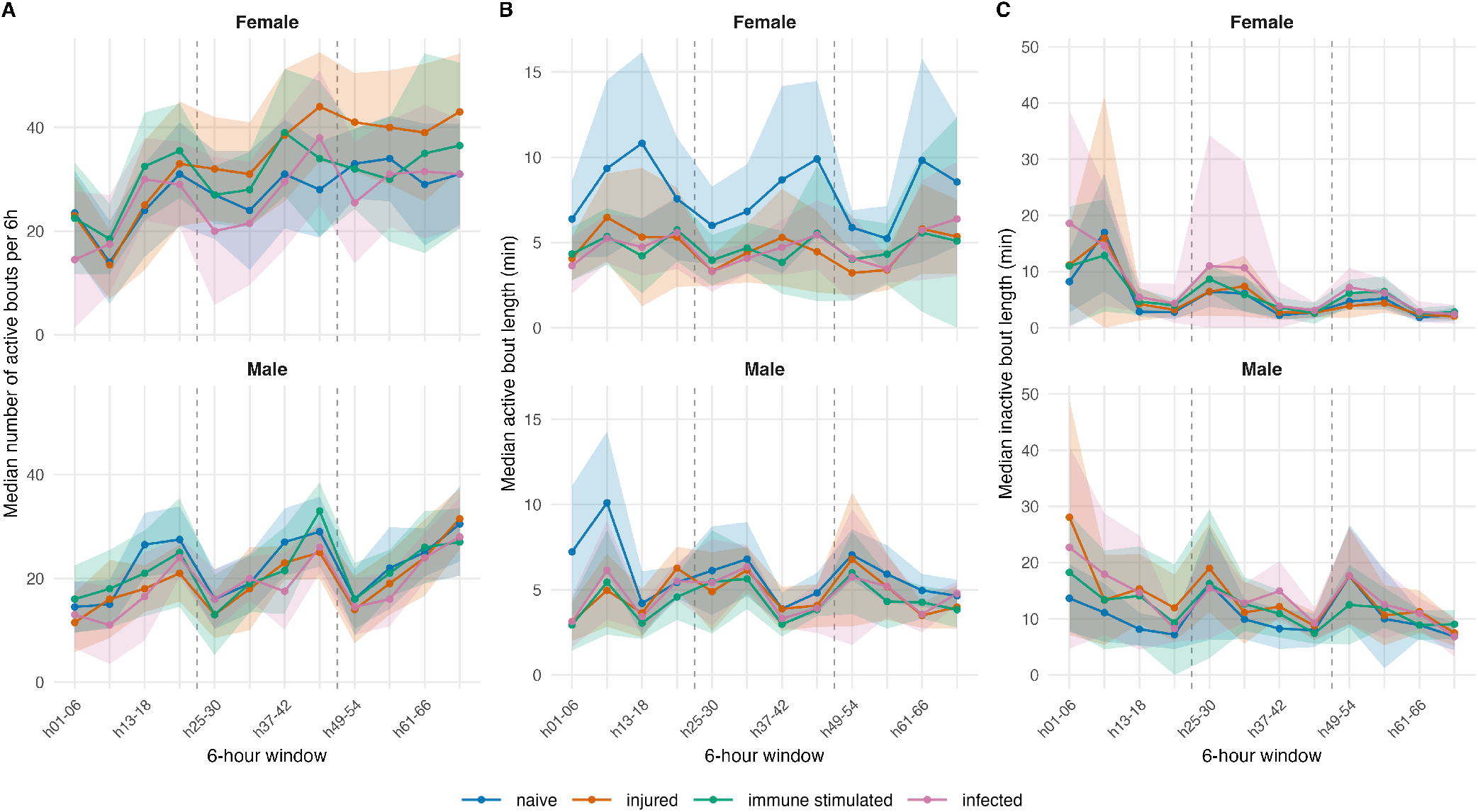
Locomotor bout structure across four treatments over the first three days post-pricking. Median values per 6-hour window for **(A)** number of active bouts per 6-hour window, active bout length (minutes), and **(C)** inactive bout length (minutes), shown separately for females (top) and males (bottom). Ribbons represent median *±* half-IQR. Dashed vertical lines indicate day boundaries (24 h and 48 h post-pricking). Naive flies (blue) showed higher bout number and longer active bouts, with shorter inactive bouts, compared to all pricking groups. Colours: naive (blue), injured (orange), immune stimulated (heat-killed *Pe* — green), infected (live *Pe* — pink).

The key differences between treatments instead emerged in the duration of active bouts rather than their number (Fig. 3B). Naive flies sustained markedly longer active bouts than all challenge groups throughout the three-day window, particularly in females (Fig. 3B; Kruskal– Wallis, females: *H* = 123, *p* < 10^−6^; males: *H* = 63.9, *p* < 10^−6^), pointing to a qualitative difference in locomotor vigour. In females, naive flies showed a median active bout length approximately 1.6-fold longer than all three challenge groups (7.35 min vs. 4.47–4.67 min); in males the effect was more modest (1.3-fold; 5.61 min vs. 4.14–4.47 min). Inactive bout length (Fig. 3C) showed a complementary, though more modest, pattern: naive flies tended to have slightly shorter resting periods than challenge groups, consistent with a higher overall drive to remain active. This effect was most apparent in males, where all three challenge groups showed median inactive bouts of 11.7–12.9 min compared to 9.84 min in naive males.

Overall, these results show that the number of periods of activity per time window was broadly similar across treatments, but active bout duration was markedly reduced by injury, immune stimulation, and live infection, particularly in females. Reduced bout length and longer rest, even in the absence of live infection, are consistent with energy conservation models of sickness behaviour and suggest host regulation of movement once activity is initiated.

### Early mortality in infected flies is linked to prolonged inactivity and brief bursts

We next inquired about the heterogeneous nature of survival times among flies infected with live *P. entomophila* (Fig. 2B). As is commonly observed in *Drosophila* infection studies, fly survival is often characterized by a sub-group of flies dying shortly after infection, while some flies are able to survive despite infection for much longer periods^36,47,48^. We therefore asked whether activity patterns could be prognostic about this heterogeneity in mortality. *P. entomophila*-infected flies were subdivided based on their total lifespan: flies that died before 12,000 minutes (200 hours) post-infection (*Infected*_*acute*_) were contrasted with those surviving beyond this threshold (*Infected*_*chronic*_), alongside naive controls (Fig. 4). *Infected*_*acute*_ flies exhibited significantly different dynamics in active bout number to *Infected*_*chronic*_ and naive flies, with this divergence being strongly sex-specific: *Infected*_*acute*_ females showed a sustained reduction in active bout number from the second 6-hour window onward, whereas *Infected*_*acute*_ males showed only a transient early reduction followed by a late reversal (Treatment × Window × Sex interaction: *F*_2,1227_ = 18.96, *p* < 0.001; Fig. 4A). Active bout length in *Infected*_*acute*_ was similarly reduced to minimal values, while inactive bout length exceeded 300 minutes per bout in females during the second day, reflecting prolonged immobility interspersed with only brief movement events (Fig. 4B–C). These bouts of extreme inactivity are consistent with a prostration-like state in moribund individuals in the hours preceding death^49^. In marked contrast, *Infected*_*chronic*_ flies maintained a bout structure relatively closer to that of naive controls, with active bout numbers, active bout lengths, and inactive bout lengths all within a relatively comparable range throughout the three-day window (Fig. 4).

**Figure 4:**
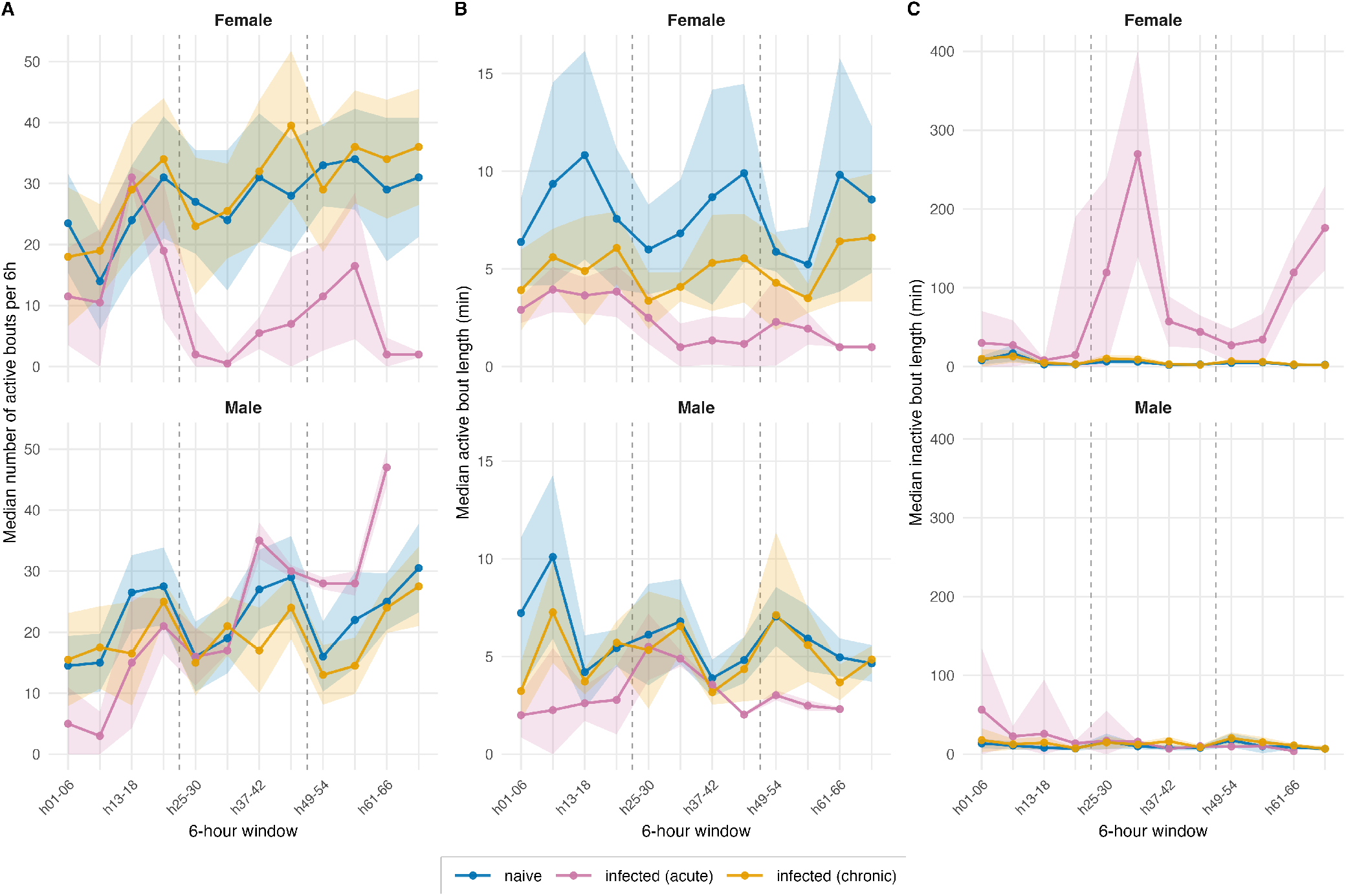
Bout structure reveals divergent behavioural trajectories within *P. entomophila*-infected flies. Median values per 6-hour window for **(A)** number of active bouts, **(B)** active bout length (minutes), and **(C)** inactive bout length (minutes), comparing naive flies with two subgroups of *P. entomophila*-infected individuals: *Infected*_*acute*_ (flies dying before 12,000 minutes post-pricking; pink) and *Infected*_*chronic*_ (flies surviving beyond 12,000 minutes; yellow). Ribbons represent median *±* half-IQR. Dashed vertical lines indicate day boundaries. *Infected*_*acute*_ flies showed a near-complete collapse of active bout structure and extreme prolongation of inactive bouts, particularly in females, while *Infected*_*chronic*_ flies maintained a bout organisation comparable to naive controls. Colours: naive (blue), *Infected*_*acute*_ (pink), *Infected*_*chronic*_ (yellow).

These results demonstrate that the behavioural response to live bacterial infection is heterogeneous within the infected population, and that early-dying individuals adopt a qualitatively distinct locomotor profile. This divergence was strongly sex-specific: females showed a sustained collapse in active bout number together with reduced active bout length and extreme prolongation of inactive bouts, whereas males showed a more transient early reduction in bout number followed by a late increase likely reflecting fragmented, ineffective movement. By contrast, infected flies that survived longer maintained a bout organisation closer to naive controls. Together, these patterns indicate that a substantial component of the behavioural change in early-dying individuals likely reflects pathology rather than an adaptive sickness behaviour.

### Bacterial infection impairs state-switching between initiation and maintenance of movement

The bout structure analysis above establishes that immune challenge affects the duration of active and inactive episodes but does not directly address the moment-to-moment action to remain in or switch between states. If sickness behaviour operates by reducing the probability of initiating movement rather than simply shortening activity bouts once started, we would expect infected flies, and particularly early-dying ones, to show an elevated probability of remaining inactive and a reduced probability of transitioning from rest to movement. Conversely, if infection also impairs the ability to sustain ongoing activity, we would additionally expect a reduced probability of remaining active and an increased probability of stopping. These two mechanisms are not mutually exclusive, but they have distinct implications: the first would manifest primarily as a failure to initiate movement, while the second would manifest as fragmented, interrupted activity bouts. Distinguishing between them requires a framework that captures both the persistence within states and the rate of switching between them.

We therefore modelled individual locomotor behaviour as a first-order two-state Markov chain, treating each minute as a discrete time step assigned to one of two states: Active (*A*_*t*_ = 1 if activity *>* 0) or Inactive (*A*_*t*_ = 0 otherwise). For each fly, a 2 × 2 transition count matrix was constructed by tabulating all consecutive minute-to-minute state pairs (*A*_*t*_, *A*_*t*+1_) over the first 24 hours. Each row was normalised by its row sum to yield a transition probability matrix, giving four probabilities: *P* (*A → A*), *P* (*A* → *I*), *P* (*I → A*), and *P* (*I → I*). Group-level matrices were obtained by summing raw counts across all flies within each treatment and sex before normalisation. The active/inactive switching ratio, defined as *P* (*I → A*)*/P* (*A* → *I*), was additionally computed per fly as a single scalar summary of the balance between activity initiation and cessation.

The resulting transition probability heatmaps revealed differences across treatment groups (Fig. 5A). The dominant transitions in all groups were state self-persistence: *P* (*A → A*) and *P* (*I → I*) were the highest probabilities in every treatment, confirming that both active and inactive states are temporally autocorrelated rather than randomly distributed. Naive flies showed the highest *P* (*A → A*) values (0.884 in females, 0.851 in males), indicating that once active, naive flies were most likely to remain so. *Infected*_*acute*_ flies displayed the lowest *P* (*A → A*) (0.785 females, 0.765 males) and the highest *P* (*A* → *I*) (0.215 females, 0.235 males) of all groups, consistent with the interpretation that early-dying individuals are unable to sustain activity episodes. *Infected*_*acute*_ flies also showed the highest *P* (*I → I*) (0.912 females, 0.960 males) and the lowest *P* (*I → A*) (0.088 females, 0.040 males), indicating that their resting periods are both more persistent and less likely to spontaneously resolve into movement. Both failure modes (impaired activity initiation and impaired activity maintenance) therefore co-occur in early-dying infected flies. *Infected*_*chronic*_ flies, by contrast, showed transition probabilities close to those of naive controls across both sexes, reinforcing the conclusion that survival trajectory is a determinant of locomotor state-switching dynamics.

**Figure 5:**
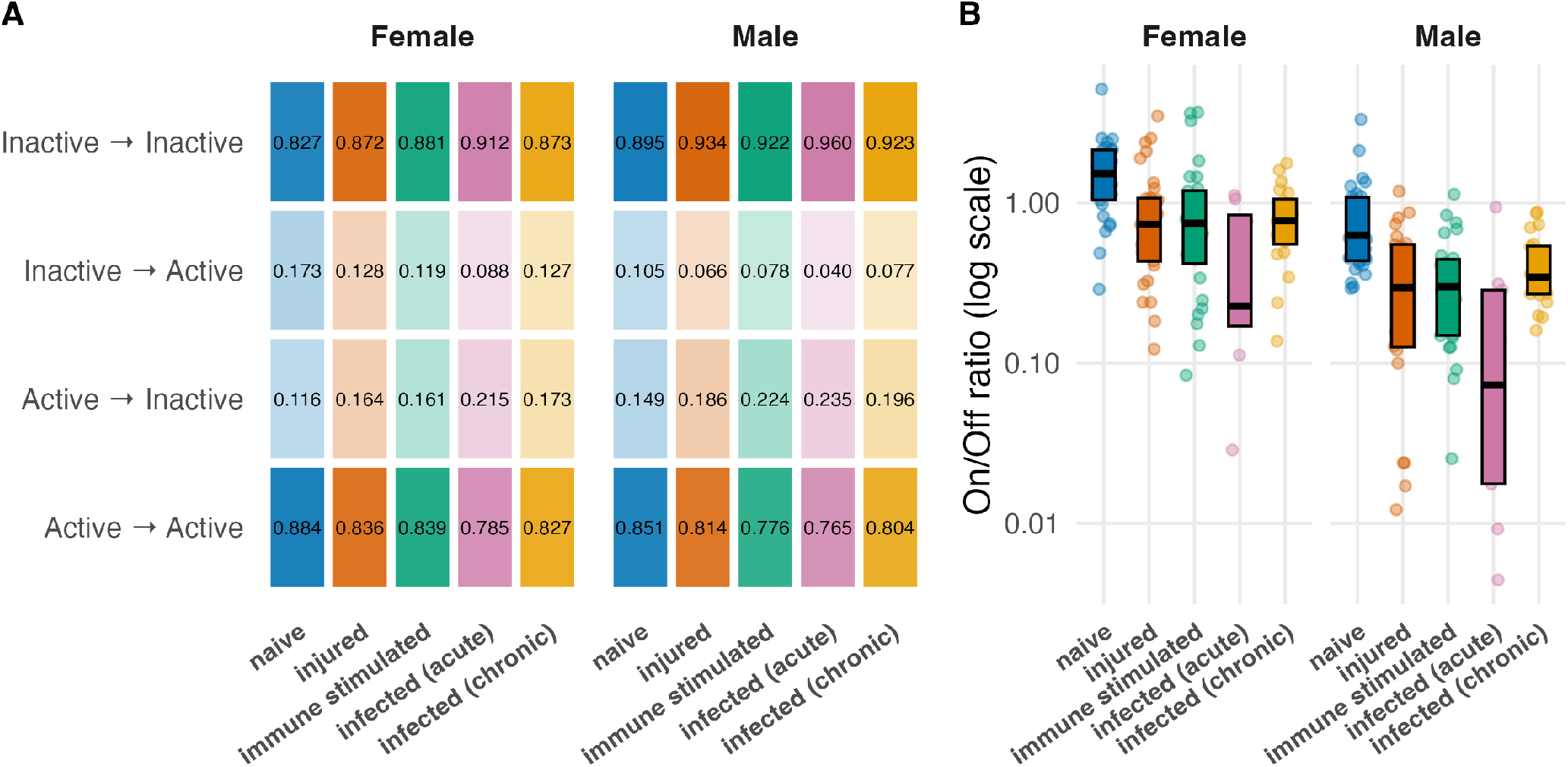
Markov chain transition probabilities and on/off switching ratio across treatment groups on Day 1 post-pricking. **(A)** Heatmaps of minute-to-minute state transition probabilities for each treatment group, shown separately for females and males. Colour intensity within each treatment is proportional to the transition probability; values are annotated in each tile. Rows correspond to the current state (Active or Inactive); columns correspond to the next state. Treatment order: naive, injured, immune stimulated, *Infected*_*acute*_, *Infected*_*chronic*_. **(B)** Per-fly on/off switching ratio (*P* (*I → A*)*/P* (*A → I*), log scale) for each treatment group and sex. Points show individual fly values (jittered); boxes show median and interquartile range. *Infected*_*acute*_ flies (pink) show a dramatically reduced on/off ratio relative to all other groups, particularly in males, reflecting a strong bias towards sustained inactivity. Colours: naive (blue), injured (orange), immune stimulated (green), *Infected*_*acute*_ (pink), *Infected*_*chronic*_ (yellow).

The per-fly active/inactive switching ratio (*P* (*I* → *A*)*/P* (*A → I*)) summarises this pattern compactly (Fig. 5B). Naive flies showed the highest median ratios in females (approximately 1.5), indicating a slight bias towards activity initiation over cessation. Injured, immune stimulated, and *Infected*_*chronic*_ flies showed lower but overlapping ratios just below unity, consistent with an almost balanced switching regime. *Infected*_*acute*_ flies were separated from all other groups, with median active/inactive ratios more than an order of magnitude lower in males (0.08) and substantially reduced in females, reflecting a strong shift towards the inactive state. The wide interquartile range in this group also indicates substantial individual variation, with flies at different stages of the dying process being captured within the same 24-hour window.

This analysis of state transitions reveals that even as early as the first day post-infection, early-dying infected flies had a reduced probability of transitioning from inactivity to activity and a reduced probability of remaining active, together with very high persistence in the inactive state. In contrast, longer-surviving infected flies had transition probabilities close to naive controls. These results demonstrate that both initiation and maintenance of movement are impaired in severe infection, consistent with a contribution of pathology and malaise beyond any adaptive reduction in voluntary activity.

### Early behavioural divergence from chronic survivors is associated with survival time in acutely dying flies

The Markov chain analysis revealed that *Infected*_*acute*_ flies exhibit a qualitatively distinct activity transition profile already on Day 1. To determine whether this early behavioural distinction was detectable at the level of raw activity levels, we quantified how strongly each *Infected*_*acute*_ fly deviated from the *Infected*_*chronic*_ activity pattern on an hour-by-hour basis across the first 24 hours post-infection (Fig. 6A). For each hour of Day 1, we expressed each *Infected*_*acute*_ fly’s activity as a Z-score relative to the *Infected*_*chronic*_ group mean and averaged the absolute deviations across all 24 hours to obtain a single divergence rate per individual (Fig. 6B).

**Figure 6:**
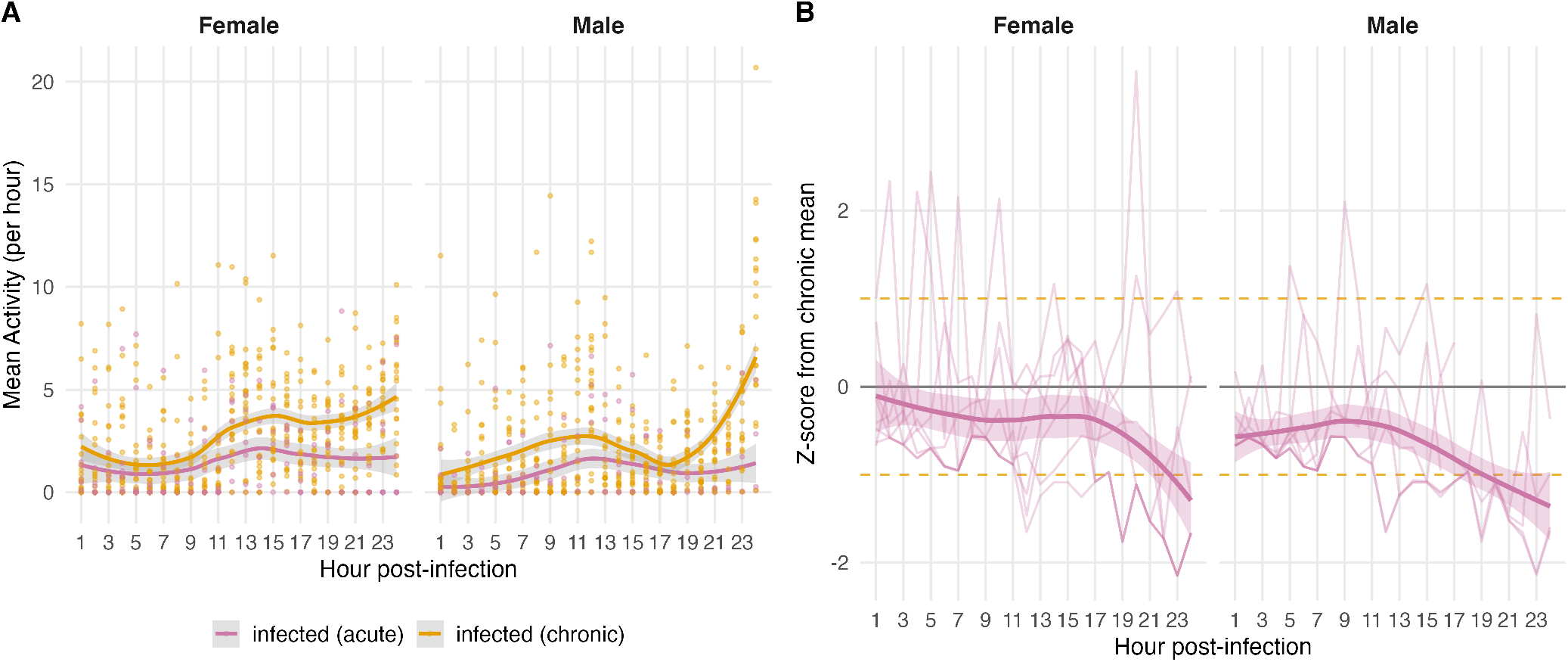
Hourly activity trajectories and Z-score deviations of *Infected*_*acute*_ flies relative to *Infected*_*chronic*_ survivors on Day 1. **(A)** Mean hourly activity for *Infected*_*acute*_ (pink) and *Infected*_*chronic*_ (orange) flies, shown separately for females and males. Lines show LOESS smooth *±* 95% CI; points are individual fly-hour means. **(B)** Per-fly Z-score deviation of *Infected*_*acute*_ activity from the *Infected*_*chronic*_ hourly mean. Each line represents one fly; the thick line and shaded band show the LOESS group trend *±* 95% CI. Dashed lines indicate the *±*1 SD threshold. Z-scores are consistently negative, indicating that acutely dying flies moved less than chronic survivors throughout Day 1.

Throughout Day 1, *Infected*_*acute*_ flies showed lower activity than chronic survivors, with Z-scores remaining below zero across the most hours in both sexes (females: 23/24 hours, males: 24/24 hours; median *Z* = − 0.57 and − 0.69 respectively; one-sample Wilcoxon test, *H*_1_: *Z* < 0; females: *p* = 2.3 × 10^−12^, males: *p* = 4.0 × 10^−18^; Fig. 6B). In females, this suppression intensified over the course of the day, as evidenced by a significant Treatment × Hour interaction (LMM: *F*_1,638_ = 15.2, *p* = 1.1 × 10^−4^); in males the interaction was not significant (*p* = 0.091), though the negative directional pattern remained consistent across all 24 hours. This indicates that acutely dying flies were already moving less than their chronic counterparts from the time of infection, and that a higher divergence rate therefore reflects a stronger suppression of activity relative to the *Infected*_*chronic*_ baseline. Males accumulated greater divergence than females across the 24-hour period (Fig. S2).

We then asked whether the degree of this early activity suppression was associated with survival time. We found a significant positive association between divergence rate and time of last movement in males (Theil–Sen trend slope = 2,574.5 min per unit Z-score, *p* = 0.019, *n* = 9): males that were suppressed more relative to chronic survivors on Day 1 tended to survive longer. The same direction of association was observed in females but was not significant (slope = 6,097.4 min per unit Z-score, *p* = 0.425, *n* = 8; Fig. S2). Given the small sample sizes inherent to the *Infected*_*acute*_ subgroup, both findings should be considered exploratory but align with the sex-specific locomotor activity patterns presented in the analyses above. This finding is consistent with short-term protective effects of behavioural restraint but could also reflect confounding by underlying damage or immune activation.

## DISCUSSION

Our goal was to distinguish adaptive, host-regulated sickness behaviours from responses to injury and pathogen-driven pathology, and to identify the behavioural features that best disentangle these drivers over time. By combining experimental partitioning of injury, sterile immune stimulation, and live bacterial infection with high resolution behavioural phenotyping, we show that (i) sickness behaviour triggered by immune challenge restructures the microstructure of locomotion by primarily shortening active bout duration and modestly prolonging rest without strongly altering bout number; (ii) live infection yields qualitatively distinct state-switching signatures only in the subset of individuals that die early, who exhibit both impaired initiation and impaired maintenance of movement; and (iii) longer-surviving infected flies maintain near normal state-transition dynamics despite being challenged by the same pathogen. These findings help resolve when reduced activity is consistent with adaptive sickness behaviour and when it signals pathology, and they provide a generalisable framework for parsing these components across systems.

### Behavioural microstructure clarifies adaptive restraint versus pathology

Across injury, immune stimulation, and live infection, we observed broadly similar bout frequencies but marked truncation of active bouts, particularly in females, coupled with modestly longer inactive bouts. This pattern fits classic models of sickness behaviour in which hosts strategically downshift energetically costly performance to buffer immune and repair investment^1,2,12^. The fact that heat killed bacteria and needle injury alone were sufficient to cause much of this restructuring indicates that a large component of the early suppression emerges from host regulation and generic damage signals, rather than requiring ongoing pathogen replication. By contrast, the collapse of bout structure in early-dying, live infected individuals was characterised by extreme prolongation of inactivity punctuated by brief, sometimes intense, movements, more likely pointing to pathology. *P. entomophila* pathogenesis includes pore-forming toxins and broad tissue damage^28–31^, which would plausibly degrade neuromuscular performance. The state transition analysis also showed that early dying flies show both a reduced probability of transitioning from rest to movement and a reduced probability of remaining active minute to minute, whereas longer surviving infected flies were indistinguishable from naive controls on these same probabilities. Thus, the behavioural features that separate adaptive restraint from pathology are less about how often flies attempt to move and more about whether they can sustain activity once begun and escape prolonged inactivity. This high-resolution discrimination of transitions between activity and inactivity may also help explain some inconsistent results in Drosophila and other taxa reporting both reduced activity and sleep induction after immune challenge^22,37,38^ and, in other contexts, increased infection induced activity^39^. Our results help reconcile these findings by showing that summary metrics (e.g., daily means) can potentially obscure qualitatively different micro-behavioural changes. A reduction in average activity can arise through shorter bouts and slightly longer rest without any change in bout initiation frequency; conversely, an apparent increase in activity could emerge from more frequent, fragmented bouts without improved bout persistence. Our data therefore underscore the value of minute scale analyses and explicit state-transition models to avoid conflating distinct behavioural mechanisms.

### Sex differences in sickness behaviour and pathology

Sexual dimorphism in immune and behavioural responses is pervasive across vertebrates and invertebrates^38,43,44^, often reflecting differences in life-history allocation and immune regulation. In flies, sex-specific coupling between metabolic state and locomotor or arousal circuits can modulate daily activity^41,42^. In this experiment, females exhibited stronger and more consistent truncation of active bouts across challenged treatments, while males showed a weaker restructuring early but a pronounced state switching collapse in the acutely dying subgroup. Two non exclusive explanations are consistent with these sex-specific behavioural patterns: (i) females may more readily enact adaptive locomotor restraint (shorter active bouts) upon immune activation, whereas (ii) males may be more vulnerable to pathological collapse of state maintenance under severe infection. Disentangling whether these patterns reflect sex-biased resistance (pathogen control) or tolerance (damage control)^15–18^ will require joint measurements of bacterial burden, damage, and behaviour within individuals.

### Heterogeneity in early behavioural profiles provide early prognostic signatures of mortality

Partitioning the infected group by subsequent survival revealed two distinct trajectories within the same challenge: a minority of acutely dying flies with a rapid behavioural collapse and a majority of chronic survivors with near normal microstructure. This echoes prior work showing that stochastic processes early in infection can tip hosts toward survival or death^47,48^. Our minute resolution Markov analysis captured divergence on Day 1, and the magnitude of early behavioural suppression in acutely dying males was positively associated with survival time. One interpretation is that, while most infected flies would eventually die, stronger early restraint modestly prolongs survival, consistent with short term protective effects of sickness behaviour. Alternatively, early suppression may simply be a consequence of host condition and variable damage, which itself varies due to stochasticity in pathogen growth dynamics. Either way, these results underline that high-frequency behavioural data may carry useful early-warning information about downstream mortality risk^36^.

### Concluding remarks

By partitioning injury, immune stimulation, and live infection, and by quantifying minute scale state switching, we show that what is often labelled “reduced activity” in infection comprises at least two mechanistically distinct phenomena. Host regulated sickness behaviour primarily reshapes the persistence of activity: shorter active bouts and modestly longer rest without large changes in bout initiation, consistent with adaptive energy reallocation. Pathogen driven pathology, in contrast, manifests as a collapse of both initiation and maintenance of movement, producing prolonged inactivity and fragmented bursts, especially in individuals fated to die early. Our framework therefore provides a tractable way to separate adaptive strategy from pathology in infection induced behavioural changes.

## MATERIALS AND METHODS

### Fly population and bacterial strain

Activity dynamics were measured in the Ashworth Advanced Outcrossed (AOx) population, a large, laboratory adapted, outbred *Drosophila melanogaster* population derived from the DGRP (Mackay et al., 2012). The AOx was established in October 2014 by setting up 100 pairwise crosses among 113 DGRP lines (Waldron, Monteith, and Vale, 2019) and has since been maintained as an outbred population on a 14 day generation cycle with a census size of 3,000–4,000 adults per generation, and incubated at 25°C under a 12:12 h light:dark cycle. *Pseudomonas entomophila* was used as the pathogen. This species infects *D. melanogaster* larvae and adults, as well as other insects, and is an aerobic, rod shaped, Gram negative bacterium with motility provided by a single polar flagellum. *P. entomophila* exhibits an optimal growth temperature of approximately 30°C and can survive between 4 and 40°C [17]. Cultures were prepared 1 day prior to assays, grown overnight with shaking, and handled under sterile conditions for the preparation of live and heat killed inocula (see Treatment groups, below).

### Treatment groups and inoculum preparation

Four treatment groups were used to distinguish the behavioural effects of injury alone, sterile immune stimulation, and live infection: (1) Naive: flies were not pricked prior to assay, providing a baseline for locomotor and sickness related behaviours without handling or injury. (2) Injured: flies were subjected to a septic injury (needle stab) using sterile phosphate buffered saline. This treatment quantifies the effect of mechanical injury and handling on sickness behaviours in the absence of microbial components. (3) Immune stimulated: this treatment measures sickness related behavioural responses attributable to sterile immune stimulation by bacterial components, without pathogen replication. Flies were subjected to a septic injury with a suspension of heat inactivated *P. entomophila*. Bacteria were grown overnight with shaking, diluted 1000x and an aliquot was heat killed in a 60–70°C water bath for 1 h. The heat inactivated suspension was then returned to shaking overnight prior to use. (4) Infected: flies were subjected to a septic injury with live *P. entomophila* prepared from the same overnight culture and standardized to an OD equivalent of 0.001 (OD=1 diluted 1000-fold). This treatment captures behavioural changes associated with active infection. For all treatments, flies were lightly anesthetized with CO2 immediately prior to the procedure, and the same injury protocol and handling were used across groups to ensure comparability. The low inoculation dose was selected to induce acute sickness while permitting survival for several days, enabling extended behavioural recording over the 16-day period.

### Activity measurement

Locomotor activity was recorded using Drosophila Activity Monitors (DAM5; Trikinetics) following established procedures [34,46]. Prior to each experiment, a 8% sucrose:2% agar solution was prepared in distilled water, autoclaved for sterilization, stored at room temperature, and re melted as needed. DAM tubes (5 mm diameter, 65 mm length) were prepared by adding approximately 1 cm of sucrose–agar medium to one end and sealing with a rubber cap to provide moisture and nutrition throughout the assay. Flies were collected upon eclosion, held on food for 2 days, anesthetized with CO2, sexed, and transferred individually into DAM tubes using a fine paint-brush. Tubes were closed with rubber caps containing a small ventilation hole [34,46]. Between 24 and 30 replicate individuals per sex and treatment were tested within a single experimental block (see Figure 1 for exact sample sizes). Each monitor included null (empty tube) or blank (no tube) channels to verify baseline signal. Activity was recorded continuously for at least 16 days (≥ 400 h) under a 12:12 h light:dark cycle at 25°C with constant temperature and humidity. Each DAM tube is bisected by four infrared beams; locomotor activity was quantified as the number of beam breaks per channel over time[34,46].

### Locomotor activity and survival analyses

To test for the effects of treatment, sex, and time on locomotor activity, we fitted a linear mixed model (LMM) using the lme4 R package. Hourly mean activity per individual was modelled as a function of treatment, sex, hour, and all two-way and three-way interactions as fixed effects, with a random intercept per individual to account for repeated measures. The model was fitted by restricted maximum likelihood (REML) using the bobyqa optimiser. Significance of fixed effects was assessed via Type III analysis of variance with Satterthwaite’s method for denominator degrees of freedom, implemented in the lmerTest package. Pairwise contrasts between treatment groups were extracted using the emmeans package, with Tukey correction for multiple comparisons across the family of four treatment levels. Contrasts were computed both averaged over the full time series and within the early 72-hour window separately by sex, to characterise sex-specific treatment effects during the acute response period. Survival was assessed from the time of pricking to the last recorded minute of activity for each individual, with flies still alive at the end of the experiment treated as right-censored observations. Kaplan-Meier survival curves were computed and visualised separately by treatment and sex. To formally test for treatment and sex effects on mortality risk, we fitted a Cox proportional hazards model using the survival R package, with treatment and sex as fixed effects and Efron’s method for handling ties. Pairwise hazard ratios between all treatment pairs were estimated on the log-hazard scale using emmeans and exponentiated to yield hazard ratios with 95% confidence intervals; p-values were adjusted using the Benjamini-Hochberg procedure.

### Bout structure analysis

To characterise the temporal organisation of locomotor activity, individual activity records were segmented into discrete bouts using run-length encoding applied to binarized activity traces (active: activity *>* 0; inactive: activity = 0). For each fly, activity was aggregated into consecutive 6-hour windows spanning the first 72 hours post-infection. Within each window, the number of active bouts, median active bout duration (min), and median inactive bout duration (min) were computed per individual. Windows containing fewer than two flies per treatment–sex combination were excluded. Group-level summaries are reported as medians with half-interquartile range (half-IQR) ribbons. To test whether treatment, sex, or their interaction affected the number of active bouts per 6-hour window, a linear mixed model was fitted with treatment, sex, and their interaction as fixed effects and a random intercept per individual to account for repeated measures across windows, using the lme4 R package. Significance of fixed effects was assessed via Type III ANOVA with Satterthwaite’s method for denominator degrees of freedom (lmerTest), and pairwise contrasts between treatment groups were extracted using emmeans with Tukey correction. To assess whether active bout duration differed between treatment groups, a Kruskal-Wallis test was applied to per-fly median active bout lengths, separately for each sex, followed by Dunn post-hoc tests with Benjamini-Hochberg correction for pairwise comparisons involving the naive group. To quantify the magnitude of differences, median fold-changes in active bout duration between naive and each challenge group were computed per sex.

### Markov chain analysis

Minute-resolution activity states (active/inactive) recorded during the first 24 hours post-infection were used to estimate first-order Markov transition probabilities for each individual. For each fly, a 2×2 transition count matrix was constructed by tallying consecutive state pairs (active→active, active→inactive, inactive→active, inactive→inactive), then row-normalised to yield transition probabilities. The on/off ratio was defined as the ratio of the inactive-to-active transition probability over the active-to-inactive transition probability, providing a dimensionless index of the tendency to initiate activity relative to the tendency to cease it. Individual-level on/off ratios are reported alongside group medians and interquartile ranges. Analyses were performed separately for females and males.

### Early behavioural divergence

To quantify the degree to which acutely infected flies (*infected*_*acute*_) diverged from the behavioural trajectory of chronically infected flies (*infected*_*chronic*_) during the first 24 hours post-infection, hourly mean activity was computed per individual. The chronic group was used as a reference, and its hourly mean (*µ*) and standard deviation (*σ*) were calculated across individuals. For each acutely infected fly, an hourly Z-score was computed as (*activity* − *µ*)*/σ*, and the absolute Z-score was taken as the instantaneous divergence. Cumulative divergence was then computed as the running sum of absolute Z-scores across hours. The hour at which each fly first exceeded a divergence threshold of one standard deviation (|*Z*| ≥ 1) was recorded as the time of first divergence. To assess whether early behavioural divergence predicted survival time, the mean hourly divergence score was regressed against time of death (hours post-infection) using Theil– Sen robust linear regression, implemented with the `mblm` package in R. Analyses were performed separately for females and males. To test whether *infected*_*acute*_ flies showed systematically lower activity than *infected*_*chronic*_ flies across Day 1, Z-scores were aggregated across all hourly observations per sex and tested against zero using a one-sided one-sample Wilcoxon signed-rank test (*H*_1_: median *Z* < 0). The proportion of hourly bins in which the median Z-score was negative was also recorded. To formally assess whether the activity difference between groups changed over the course of Day 1, a linear mixed model (LMM) was fitted with hourly mean activity as the response variable, Treatment, Hour, and their interaction as fixed effects, and individual fly as a random intercept: Activity *∼* Treatment × Hour+(1 | FlyID) using the `lmerTest` package in R with a `bobyqa` optimiser. Type III ANOVAs were used to evaluate fixed effects. Additionally, per-hour pairwise comparisons between *infected*_*acute*_ and *infected*_*chronic*_ were performed using two-sample Wilcoxon tests, with *p*-values corrected for multiple comparisons using the Benjamini–Hochberg procedure. All analyses were performed separately for females and males.

## RESOURCE AVAILABILITY

### Lead contact

Requests for further information and resources should be directed to and will be fulfilled by Alejandro V Cano and Pedro Vale.

### Materials availability

This study did not generate new materials.

### Data and code availability

The data and code used for the analyses and figures of this paper is available at the GitHub repository https://github.com/alejvcano/flies_sickness_behaviour.

## ACKNOWLEDGMENTS

This work was funded by a Royal Society International Exchange grant IES\R2\222092 awarded to PFV and VD, and a School of Biological Sciences Seed Fund (University of Edinburgh) awarded to PFV. PFV also acknowledges funding support from the BBSRC (Award UKRI709). For the purpose of open access, the authors have applied a Creative Commons Attribution (CC BY) licence to any Author Accepted Manuscript version arising from this submission.

## AUTHOR CONTRIBUTIONS

Conceptualization, A.V.C., D.N., K.M., V.D., and P.V.; methodology, A.V.C., D.N., K.M., V.D., and P.V.; investigation, A.V.C., D.N. and K.M.; writing-–original draft, A.V.C. and P.V.; writing-–review & editing, A.V.C., D.N., K.M., V.D., and P.V.; funding acquisition, V.D. and P.V.; resources, V.D. and P.V.; supervision, V.D. and P.V.

---

**Figure S1:**
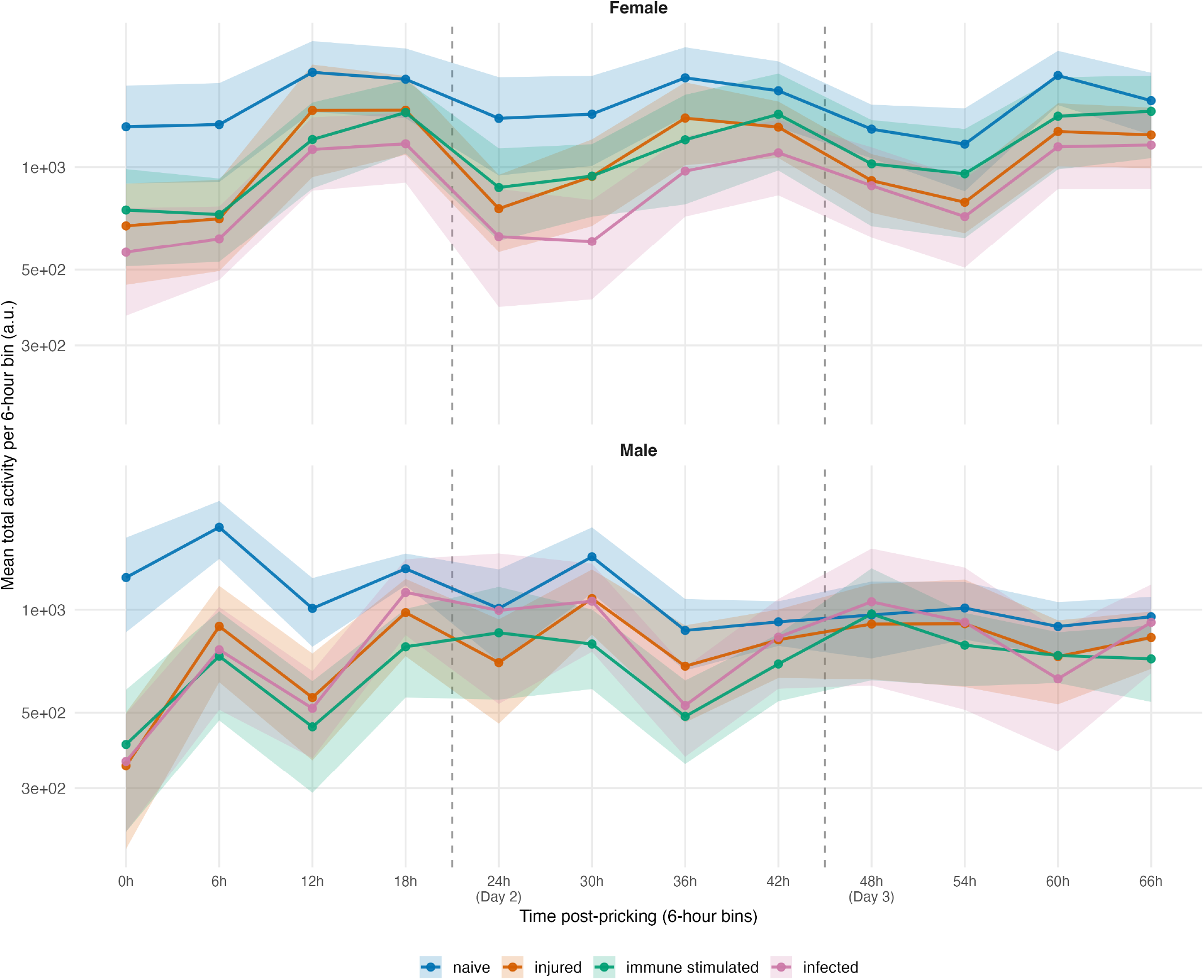
Mean locomotor activity in 6-hour bins over the first three days post-injection. Mean total activity per 6-hour bin (arbitrary units, log scale) for each treatment group, shown separately for females (top) and males (bottom). Points represent group means; shaded ribbons indicate 95% confidence intervals. Dashed vertical lines mark transitions between Days 1, 2, and 3. Naive flies (blue) maintained consistently higher activity than all other groups across both sexes. Injured flies (orange) showed activity comparable to naive controls. Immune stimulated (green) and infected flies (pink) displayed lower activity from the earliest bins, with the suppression more pronounced in males. The oscillatory pattern within each day reflects the underlying circadian rhythm, with peaks in the 6 h and 18 h bins corresponding to morning and evening activity bouts. Colours: naive (blue), injured (orange), immune stimulated (green), *P. entomophila* (pink).

**Figure S2:**
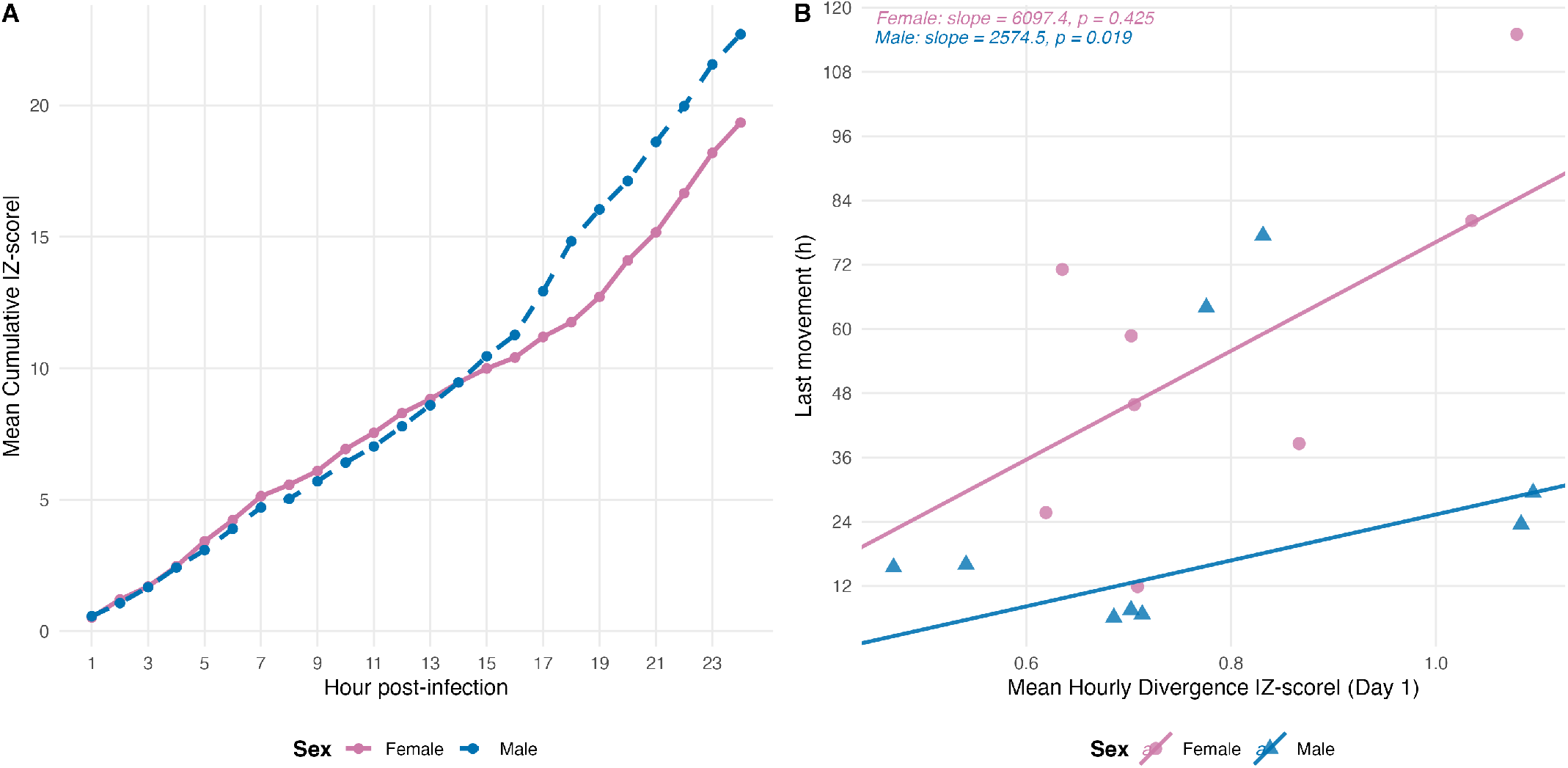
Day 1 behavioural divergence from chronic survivors in acutely dying flies. **(A)** Mean cumulative absolute Z-score deviation per hour across Day 1 for *Infected*_*acute*_ females (pink, solid) and males (blue, dashed). **(B)** Mean hourly divergence rate on Day 1 versus time of last movement for individual *Infected*_*acute*_ flies (females: circles; males: triangles). Lines show Theil–Sen trends; slope and *p*-values are indicated for each sex.

## References

1. Hart, B.L. (1988). Biological basis of the behavior of sick animals. Neuroscience and Biobehavioral Reviews 12, 123–137. doi: 10.1016/s0149-7634(88)80004-6.

2. Hart, B.L. (2011). Behavioural defences in animals against pathogens and parasites: parallels with the pillars of medicine in humans. Phil. Trans. R. Soc. B 366, 3406–3417. URL: http://rstb.royalsocietypublishing.org/content/366/1583/3406. doi: 10.1098/rstb.2011.0092.

3. Dantzer, R., and Kelley, K.W. (2007). Twenty years of research on cytokine-induced sickness behavior. Brain, Behavior, and Immunity 21, 153–160. URL: http://www.sciencedirect.com/science/article/pii/S088915910600300X. doi: 10.1016/j.bbi.2006.09.006.

4. Lopes, P.C., French, S.S., Woodhams, D.C., and Binning, S.A. (2021). Sickness behaviors across vertebrate taxa: proximate and ultimate mechanisms. Journal of Experimental Biology 224, jeb225847. URL:. doi: 10.1242/jeb.225847.

5. Vale, P.F. (2018). Disease Tolerance: Linking Sickness Behaviours to Metabolism Helps Mitigate Malaria. Current Biology 28, R606–R607. URL: https://drive.google.com/file/d/1QuW7LWh0PpTuPEd94yZ_aZ1WfUM5YXSC/view?usp=sharing. doi: 10.1016/j.cub.2018.04.031.

6. Sullivan, K., Fairn, E., and Adamo, S.A. (2016). Sickness behaviour in the cricket Gryllus texensis: Comparison with animals across phyla. Behavioural Processes 128, 134–143. doi: 10.1016/j.beproc.2016.05.004.

7. Kazlauskas, N., Klappenbach, M., Depino, A.M., and Locatelli, F.F. (2016). Sickness Behavior in Honey Bees. Frontiers in Physiology 7. URL: http://www.ncbi.nlm.nih.gov/pmc/articles/PMC4924483/. doi: 10.3389/fphys.2016.00261.

8. Milutinović, B., and Schmitt, T. (2022). Chemical cues in disease recognition and their immunomodulatory role in insects. Current Opinion in Insect Science 50, 100884. URL: https://www.sciencedirect.com/science/article/pii/S2214574522000190. doi: 10.1016/j.cois.2022.100884.

9. Adamo, S.A. (2006). Comparative psychoneuroimmunology: evidence from the insects. Behavioral and Cognitive Neuroscience Reviews 5, 128–140. doi: 10.1177/1534582306289580.

10. Lopes, P.C. (2014). When is it socially acceptable to feel sick? Proceedings of the Royal Society of London B: Biological Sciences 281, 20140218. URL: http://rspb.royalsocietypublishing.org/content/281/1788/20140218. doi: 10.1098/rspb.2014.0218.

11. Ezenwa, V.O., Archie, E.A., Craft, M.E., Hawley, D.M., Martin, L.B., Moore, J., and White, L. (2016). Host behaviour–parasite feedback: an essential link between animal behaviour and disease ecology. Proc. R. Soc. B 283, 20153078. URL: http://rspb.royalsocietypublishing.org/content/283/1828/20153078. doi: 10.1098/rspb.2015.3078.

12. Shakhar, K., and Shakhar, G. (2015). Why Do We Feel Sick When Infected—Can Altruism Play a Role? PLOS Biology 13, e1002276. URL: https://journals.plos.org/plosbiology/article?id=10.1371/journal.pbio.1002276. doi: 10.1371/journal.pbio.1002276.

13. Stockmaier, S., Bolnick, D.I., Page, R.A., and Carter, G.G. (2020). Sickness effects on social interactions depend on the type of behaviour and relationship. Journal of Animal Ecology 89, 1387–1394. URL: https://onlinelibrary.wiley.com/doi/abs/10.1111/1365-2656.13193. doi: 10.1111/1365-2656.13193. eprint: https://besjournals.onlinelibrary.wiley.com/doi/pdf/10.1111/1365-2656.13193.

14. Lopes, P.C., Wingfield, J.C., and Bentley, G.E. (2012). Lipopolysaccharide injection induces rapid decrease of hypothalamic GnRH mRNA and peptide, but does not affect GnIH in zebra finches. Hormones and Behavior 62, 173–179. URL: https://www.sciencedirect.com/science/article/pii/S0018506X12001729. doi: 10.1016/j.yhbeh.2012.06.007.

15. Cumnock, K., Gupta, A.S., Lissner, M., Chevee, V., Davis, N.M., and Schneider, D.S. (2018). Host Energy Source Is Important for Disease Tolerance to Malaria. Current biology: CB 28, 1635–1642.e3. doi: 10.1016/j.cub.2018.04.009.

16. Ganeshan, K., Nikkanen, J., Man, K., Leong, Y.A., Sogawa, Y., Maschek, J.A., Van Ry, T., Chagwedera, D.N., Cox, J.E., and Chawla, A. (2019). Energetic Trade-Offs and Hypometabolic States Promote Disease Tolerance. Cell 177, 399–413.e12. URL: http://www.sciencedirect.com/science/article/pii/S0092867419301138. doi: 10.1016/j.cell.2019.01.050.

17. Lopes, P.C., Springthorpe, D., and Bentley, G.E. (2014). Increased activity correlates with reduced ability to mount immune defenses to endotoxin in zebra finches. Journal of Experimental Zoology Part A: Ecological Genetics and Physiology 321, 422–431. URL: https://onlinelibrary.wiley.com/doi/abs/10.1002/jez.1873. doi: 10.1002/jez.1873. eprint: https://onlinelibrary.wiley.com/doi/pdf/10.1002/jez.1873.

18. Prakash, A., Monteith, K.M., and Vale, P.F. (2022). Mechanisms of damage prevention, signalling, and repair impact disease tolerance. Proceedings of the Royal Society B pp. 2021.10.03.462916. URL: https://www.biorxiv.org/content/10.1101/2021.10.03.462916v2. doi: 10.1101/2021.10.03.462916.

19. Soares, M.P., Gozzelino, R., and Weis, S. (2014). Tissue damage control in disease tolerance. Trends in Immunology 35, 483–494. URL: http://www.sciencedirect.com/science/article/pii/S147149061400132X. doi: 10.1016/j.it.2014.08.001.

20. Lopes, P.C., Block, P., and König, B. (2016). Infection-induced behavioural changes reduce connectivity and the potential for disease spread in wild mice contact networks. Scientific Reports 6, 31790. URL: http://www.nature.com/srep/2016/160822/srep31790/full/srep31790.html. doi: 10.1038/srep31790.

21. Romano, V., Lussiana, A., Monteith, K., MacIntosh, A.J.J., and Vale, P. (2022). Host and pathogen drivers of infection-induced changes in social aggregation behaviour. bioRxiv.

22. Siva-Jothy, J.A., and Vale, P.F. (2019). Viral infection causes sex-specific changes in fruit fly social aggregation behaviour. Biology Letters 15, 20190344. URL: https://royalsocietypublishing.org/doi/10.1098/rsbl.2019.0344. doi: 10.1098/rsbl.2019.0344.

23. Shoemaker, K.L., Parsons, N.M., and Adamo, S.A. (2006). Egg-laying behaviour following infection in the cricket Gryllus texensis. Canadian Journal of Zoology 84, 412–418. URL: http://www.nrcresearchpress.com/doi/10.1139/z06-013. doi: 10.1139/z06-013.

24. Hudson, A.L., Moatt, J.P., and Vale, P.F. (2019). Terminal investment strategies following infection are dependent on diet. Journal of Evolutionary Biology n/a. URL: https://onlinelibrary.wiley.com/doi/abs/10.1111/jeb.13566. doi: 10.1111/jeb.13566.

25. Westlake, H., Hanson, M.A., and Lemaitre, B. (2024). The Drosophila immunity handbook. First ed.. EPFL Press. ISBN 978-2-8323-2267-3. URL: https://www.epflpress.org/produit/1514/9782889156467. doi: 10.55430/6304TDIHVA01.

26. Dubnau, J. (2014). Behavioral Genetics of the Fly (Drosophila Melanogaster). Cambridge University Press. ISBN 978-1-107-00903-5.

27. Aranha, M.M., and Vasconcelos, M.L. (2018). Deciphering Drosophila female innate behaviors. Current Opinion in Neurobiology 52, 139–148. URL: https://www.sciencedirect.com/science/article/pii/S0959438818300473. doi: 10.1016/j.conb.2018.06.005.

28. Opota, O., Vallet-Gély, I., Vincentelli, R., Kellenberger, C., Iacovache, I., Gonzalez, M.R., Roussel, A., van der Goot, F.G., and Lemaitre, B. (2011). Monalysin, a novel ß-pore-forming toxin from the drosophila pathogen pseudomonas entomophila, contributes to host intestinal damage and lethality. PLoS Pathogens 7. doi: 10.1371/journal.ppat.1002259.

29. Vodovar, N., Vinals, M., Liehl, P., Basset, A., Degrouard, J., Spellman, P., Boccard, F., and Lemaitre, B. (2005). Drosophila host defense after oral infection by an entomopathogenic Pseudomonas species. Proceedings of the National Academy of Sciences of the United States of America 102, 11414–9. URL: http://www.pnas.org/content/102/32/11414. doi: 10.1073/pnas.0502240102.

30. Liehl, P., Blight, M., Vodovar, N., Boccard, F., and Lemaitre, B. (2006). Prevalence of Local Immune Response against Oral Infection in a Drosophila/Pseudomonas Infection Model. PLoS Pathog 2, e56. URL: http://dx.plos.org/10.1371/journal.ppat.0020056. doi: 10.1371/journal.ppat.0020056.

31. Prakash, A., Bonnet, M., Monteith, K.M., and Vale, P.F. (2021). The Jak/Stat pathway mediates disease tolerance during systemic bacterial infection in Drosophila. bioRxiv. URL: https://www.biorxiv.org/content/10.1101/2021.09.23.461578v1. doi: 10.1101/2021.09.23.461578. Company: Cold Spring Harbor Laboratory Distributor: Cold Spring Harbor Laboratory Label: Cold Spring Harbor Laboratory Type: article.

32. Prakash, A., Monteith, K.M., and Vale, P.F. (2023). Negative immune regulation contributes to disease tolerance in Drosophila. bioRxiv pp. 2021.09.23.461574. URL: https://www.biorxiv.org/content/10.1101/2021.09.23.461574v3. doi: 10.1101/2021.09.23.461574.

33. Shaw, P.J., Cirelli, C., Greenspan, R.J., and Tononi, G. (2000). Correlates of sleep and waking in Drosophila melanogaster. Science 287, 1834–7. doi: 10.1126/science.287.5459.1834. Number: 5459.

34. Chiu, J.C., Low, K.H., Pike, D.H., Yildirim, E., and Edery, I. (2010). Assaying Locomotor Activity to Study Circadian Rhythms and Sleep Parameters in <em>Drosophila</em>. Journal of Visualized Experiments. URL: http://www.jove.com/video/2157/assaying-locomotor-activity-to-study-circadian-rhythms-sleep. doi: 10.3791/2157.

35. Anderson, L., Camus, M.F., Monteith, K.M., Salminen, T.S., and Vale, P.F. (2022). Variation in mitochondrial DNA affects locomotor activity and sleep in Drosophila melanogaster. Heredity pp. in press. URL: https://www.biorxiv.org/content/10.1101/2021.10.30.464953v1. doi: 10.1101/2021.10.30.464953.

36. Kutzer, M.A.M., Abdullateef, S., Cano, A.V., Soare-Nguyen, I.L., Escudero, J., Dakos, V., and Vale, P.F. (2025). Detecting infection-related mortality using dynamical statistical indicators of high-resolution activity time series. BioRxiv.

37. Kuo, T.H., Pike, D.H., Beizaeipour, Z., and Williams, J.A. (2010). Sleep triggered by an immune response in Drosophila is regulated by the circadian clock and requires the NFkappaB Relish. BMC Neuroscience 11, 17. URL: http://www.biomedcentral.com/1471-2202/11/17. doi: 10.1186/1471-2202-11-17.

38. Vale, P.F., and Jardine, M.D. (2015). Sex-specific behavioural symptoms of viral gut infection and Wolbachia in Drosophila melanogaster. Journal of Insect Physiology 82, 28–32. doi: 10.1016/j.jinsphys.2015.08.005.

39. Vincent, C.M., Beckwith, E.J., Simoes da Silva, C.J., Pearson, W.H., Kierdorf, K., Gilestro, G.F., and Dionne, M.S. (2022). Infection increases activity via Toll dependent and independent mechanisms in Drosophila melanogaster. PLOS Pathogens 18, e1010826– e1010826. URL: https://dx.plos.org/10.1371/journal.ppat.1010826. doi: 10.1371/journal.ppat.1010826.

40. Gordon, J., and Masek, P. (2021). Excessive energy expenditure due to acute physical restraint disrupts Drosophila motivational feeding response. Scientific Reports 11, 24208. URL: https://pmc.ncbi.nlm.nih.gov/articles/PMC8683507/. doi: 10.1038/s41598-021-03575-3.

41. Lee, S.H., Cho, E., Yoon, S.E., Kim, Y., and Kim, E.Y. (2021). Metabolic control of daily locomotor activity mediated by tachykinin in Drosophila. Communications Biology 4, 693. URL: https://pmc.ncbi.nlm.nih.gov/articles/PMC8184744/. doi: 10.1038/s42003-021-02219-6.

42. Shafer, O.T., and Keene, A.C. (2021). The Regulation of Drosophila Sleep. Current Biology 31, R38–R49. URL: https://www.cell.com/current-biology/abstract/S0960-9822(20)31659-6. doi: 10.1016/j.cub.2020.10.082.

43. Klein, S.L., and Flanagan, K.L. (2016). Sex differences in immune responses. Nature Reviews Immunology 16, 626–638. URL: http://www.nature.com/nri/journal/v16/n10/full/nri.2016.90.html. doi: 10.1038/nri.2016.90.

44. Belmonte, R.L., Corbally, M.K., Duneau, D.F., and Regan, J.C. (2019). Sexual Dimorphisms in Innate Immunity and Responses to Infection in Drosophila melanogaster. Frontiers in Immunology 10, 3075. doi: 10.3389/fimmu.2019.03075.

45. Kutzer, M.A.M., Cornish, B., Jamieson, M., Zawistowska, O., Monteith, K.M., and Vale, P.F. (2024). Mitochondrial background can explain variable costs of immune deployment. Journal of Evolutionary Biology. URL: https://www.biorxiv.org/content/10.1101/2023.10.04.560830v1. doi: 10.1101/2023.10.04.560830.

46. Pfeiffenberger, C., Lear, B.C., Keegan, K.P., and Allada, R. (2010). Locomotor Activity Level Monitoring Using the Drosophila Activity Monitoring (DAM) System. Cold Spring Harbor Protocols 2010, pdb.prot5518. URL: http://cshprotocols.cshlp.org/content/2010/11/pdb.prot5518. doi: 10.1101/pdb.prot5518.

47. Duneau, D., Ferdy, J.B., Revah, J., Kondolf, H., Ortiz, G.A., Lazzaro, B.P., and Buchon, N. (2017). Stochastic variation in the initial phase of bacterial infection predicts the probability of survival in D. melanogaster. eLife 6, e28298. URL:. doi: 10.7554/eLife.28298.

48. Chen, J., Lin, G., Ma, K., Li, Z., Liégeois, S., and Ferrandon, D. (2024). A specific innate immune response silences the virulence of Pseudomonas aeruginosa in a latent infection model in the Drosophila melanogaster host. PLoS pathogens 20, e1012252. doi: 10.1371/journal.ppat.1012252.

49. Ijaopo, E.O., Zaw, K.M., Ijaopo, R.O., and Khawand-Azoulai, M. (2023). A Review of Clinical Signs and Symptoms of Imminent End-of-Life in Individuals With Advanced Illness. Gerontology and Geriatric Medicine 9, 23337214231183243. URL: https://pmc.ncbi.nlm.nih.gov/articles/PMC10327414/. doi: 10.1177/23337214231183243.

